# Covalent remodeling of CRBN creates a non-canonical neosubstrate interface with NTAQ1

**DOI:** 10.64898/2026.07.14.738385

**Authors:** Andres H. de la Peña, Justin T. Cruite, Jianwei Che, Mary E. Matyskiela, Philip P. Chamberlain, Eric S. Fischer, Lyn H. Jones

## Abstract

Molecular glue degrader EM12-FS covalently modifies cereblon (CRBN) His353, enabling selective recruitment of the neosubstrate NTAQ1 to the CRL4^CRBN^ ubiquitin ligase. We determined the cryo-EM structure of the NTAQ1–EM12-FS–CRBN–DDB1 complex, revealing a non-canonical neosubstrate interface created by covalent remodeling of the CRBN sensor loop. Imidazylation repositions His353 to eliminate the steric clash that prevents NTAQ1 engagement by reversible IMiDs, and the engineered interface is stabilized by a distinctive T-shaped C-H/π interaction between sulfated His353 and NTAQ1 Phe126. Biochemical and mutational analyses define the determinants of ternary complex formation and ubiquitination. These findings show that site-specific synthetic modification of CRBN can reprogram induced-proximity pharmacology, expanding specificity beyond the G-loop degron and establishing a framework for covalent engineering of new degrader modalities.

## Introduction

Immunomodulatory imide drugs (IMiDs) thalidomide, lenalidomide and pomalidomide bind cereblon (CRBN),^1^ the substrate adaptor of the E3 ubiquitin ligase complex DDB1–Cul4–Rbx1–CRBN (CRL4^CRBN^).^2,3^ IMiD binding remodels the CRBN surface to recruit neosubstrates for polyubiquitination and proteasomal degradation. Most known neosubstrates share a β-hairpin loop containing a critical glycine that enables engagement with the IMiD– CRBN surface. This canonical “G-loop degron” is a defining structural feature of CRBN neosubstrates,^4,5^ many of which were previously considered undruggable due to the absence of conventional ligandable pockets.^6^ Clinically approved IMiDs degrade IKZF1/3 in multiple myeloma and CK1α in del(5q) myelodysplastic syndrome,^7^ and additional therapeutic candidates targeting other neosubstrates are emerging. Although many proteins contain G-loop motifs,^4,6^ the therapeutic landscape beyond this degron remains largely unexplored.^8-11^

We reasoned that site-specific synthetic manipulation of protein surfaces could mimic post-translational modifications or hotspot mutations that rewire protein interactions and generate new functional states.^12^ Covalent engagement of E3 ligases offers a route to impose novel physicochemical features and enforce conformations inaccessible to reversible ligands, enabling recruitment of non-canonical substrates.^12-14^ This strategy also offers a pharmacological advantage: irreversible modification can decouple pharmacokinetics from pharmacodynamics by locking the ligase into a new specificity state even after compound clearance. Although cysteine-targeting electrophiles dominate covalent drug discovery and chemical biology, the thalidomide-binding domain (TBD) of CRBN contains no suitably positioned cysteine for covalent engagement. However, a histidine residue (His353) lies at the tip of the ‘sensor loop’ adjacent to reversible binding IMiDs and contributes to neosubstrate engagement,^15^ making it an attractive alternative nucleophile for covalent neofunctionalization.

We previously developed covalent molecular glues based on the IMiD derivative EM12, incorporating sulfonyl exchange electrophiles to react with His353.^13^ Functionalization at the EM12 6-position yielded the sulfonyl fluoride EM12-SF and fluorosulfate EM12-FS (Fig. 1A), both of which site-specifically modify His353, constituting the first rational application of sulfonyl exchange chemistry to target a histidine residue. Computational modeling predicted that covalent modification would reposition His353 and introduce steric clashes with the G-loop degron, thereby preventing recruitment of canonical neosubstrates. Consistent with this prediction, EM12-SF did not induce detectable degradation in cells, whereas EM12-FS exclusively degraded *N*-terminal glutamine amidase 1 (NTAQ1), a previously unrecognized CRBN neosubstrate that initiates the Arg/N-degron pathway.^16^ Covalent engagement of His353 was required for EM12-FS-dependent NTAQ1 recruitment and degradation.^13^ These observations suggested that covalent remodeling of CRBN could create new ligandable surfaces and enable recruitment of substrates inaccessible to reversible IMiDs.

**Fig. 1.**
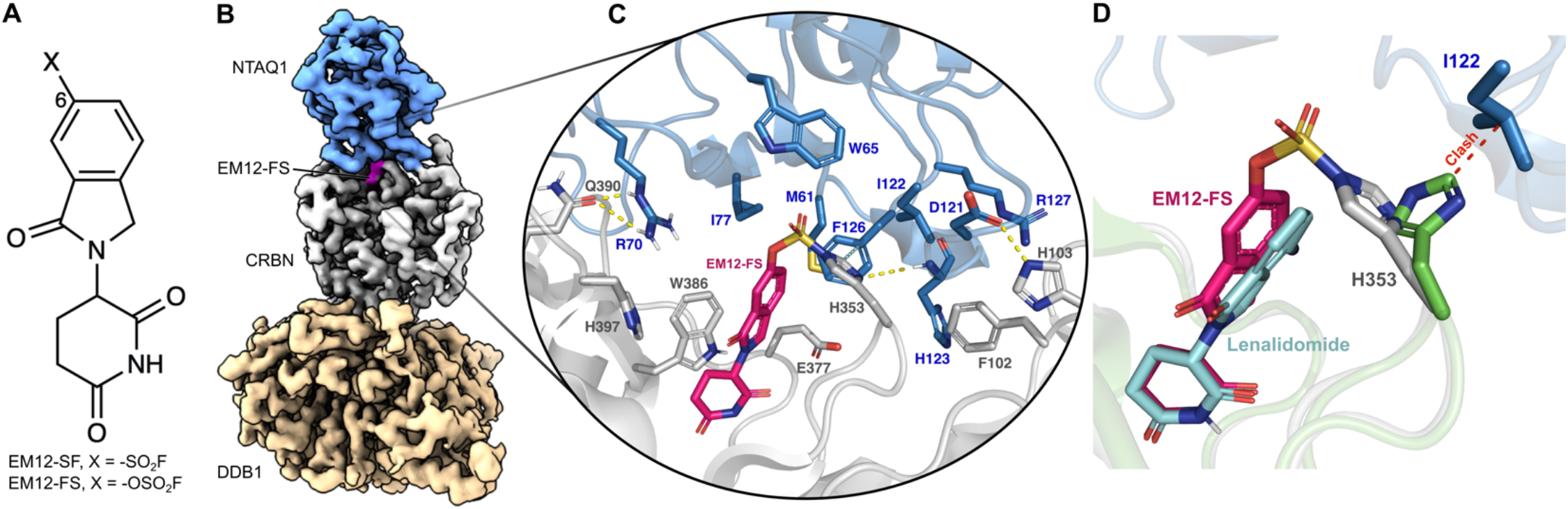
Cryo-EM structure of the NTAQ1–EM12-FS–CRBN–DDB1 complex. **A)** Chemical structure of covalent CRBN modulators EM12-SF and EM12-FS. **B**) Experimental density of the ternary complex depicting NTAQ1 (blue), EM12-FS (magenta), CRBN (gray), and DDB1 (beige). **C)** Detailed view of the EM12-FS interface showing hydrogen-bonding interactions (yellow dotted lines). **D)** Binding mode comparison of EM12-FS and lenalidomide (cyan, PDB 5FQD)^4^ showing the clash (red dotted line) between His353 (green) in the lenalidomide structure and Ile122 in NTAQ1.

## Results

To define the molecular basis of this selectivity, we determined the structure of the NTAQ1–EM12-FS–CRBN– DDB1 complex using cryo-EM to a resolution of 3.2 Å at the ternary interface (Figs. 1B, 1C, Fig. S1 and Table 1). The reconstruction reveals that covalent engagement of His353 generates a new ligandable surface distinct from all reversible IMiD complexes. To ensure that the observed conformation reflects this engineered state, CRBN was covalently pre-labeled with EM12-FS prior to ternary complex assembly.

**Table 1:** Cryo-EM data collection, refinement, and validation statistics.

| CRBN/DDB1, EM12-FS, NTAQ1 Ternary complex<br>PDB: 36MP<br>EMDB: EMD-77682 |  |
| --- | --- |
| **Data collection and processing** |  |
| Magnification | 81,000x |
| Voltage (kV) | 300 |
| Electron exposure (e-/Å<sup>2</sup>) | 41 |
| Defocus range (μm) | -0.8 to -2.0 |
| Pixel size (Å) | 1.072 |
| Symmetry imposed | C<sub>1</sub> |
| Initial particle images (no.) | 4,481,872 |
| Final particle images (no.) | 109,337 |
| Map resolution (Å) | 3.3 |
| FSC threshold | 0.143 |
| Map resolution range (Å) | 3.2 to 4.8 |
| **Refinement** |  |
| Initial model used (AF Code) | Q16531-F1, Q96SW2-F1, Q96HA8-F1 |
| Model resolution (Å) | 3.3 |
| FSC threshold | 0.143 |
| Model resolution range (Å) | 3.2 to 4.8 |
| Map sharpening *B* factor (Å<sup>2</sup>) | -80.43 |
| **Model composition** |  |
| Non-hydrogen atoms | 10,847 |
| Protein residues | 1375 |
| Ligands | UNK |
| ***B* factors (Å<sup>2</sup>, mean)** |  |
| Protein | 71.86 |
| Ligand | 33.62 |
| **R.m.s. deviations** |  |
| Bond lengths (Å) | 0.006 |
| Bond angles (°) | 1.002 |
| **Validation** |  |
| MolProbity score | 1.49 |
| Clashscore | 3.92 |
| Poor rotamers (%) | 0.50 |
| **Ramachandran plot** |  |
| Favored (%) | 95.51 |
| Allowed (%) | 4.49 |
| Disallowed (%) | 0 |

As expected, the EM12-FS glutarimide binds the tri-tryptophan cage of the TBD and the fluorosulfate forms an imidazylate with His353. NTAQ1 engages the EM12-FS–CRBN adduct primarily through residues at the periphery of its α-β-α amidohydrolase fold (Fig. S2). A defining feature of the interface is a T-shaped edge-to-face π-interaction between the polarized C-H at the 2-position of sulfated His353 and the aromatic ring of NTAQ1 Phe126. Additional complementarity arises from hydrogen bonds and hydrophobic contacts outside the IMiD pocket (Fig. 1C and Table S1).

Sulfation repositions the His353 imidazole to avoid a steric clash with NTAQ1 Ile122 (Fig. 1D, Fig. S3), explaining why NTAQ1 is not recruited by reversible binding IMiDs. This structure captures a synthetically engineered CRBN specificity state and demonstrates that covalent modification can impose conformations inaccessible to reversible ligands.

Although EM12-SF differs from EM12-FS by only a single oxygen atom, the resulting change in electrophile geometry alters the orientation of His353 and disrupts key hydrophobic contacts with Ile122 and Phe126 (Fig. S4). The cryo-EM structure and MM-GBSA calculations together show that productive ternary complex formation requires precise warhead-imposed positioning of His353. These data establish a potential design principle for covalent molecular glues: the electrophile must not only react but must enforce a neofunctionalized CRBN conformation compatible with the intended substrate.^13^

Mutational analysis of NTAQ1 confirmed that the majority of the hydrophobic residues interacting with EM12-FS are indispensable for ternary complex formation (TCF) (Fig. 2, Figs. S5-S7). Additional residues at the protein-protein interface (PPI) modulated complex stability without directly contacting the compound, decreasing (D120, D121) or increasing (Q125, R157) TCF (Fig. 2, Figs. S6, S7). Consistent with the cryo-EM structure, the F126 residue is essential for both TCF and NTAQ1 ubiquitination (Fig. 2, Figs. S7, S8).

**Fig. 2.**
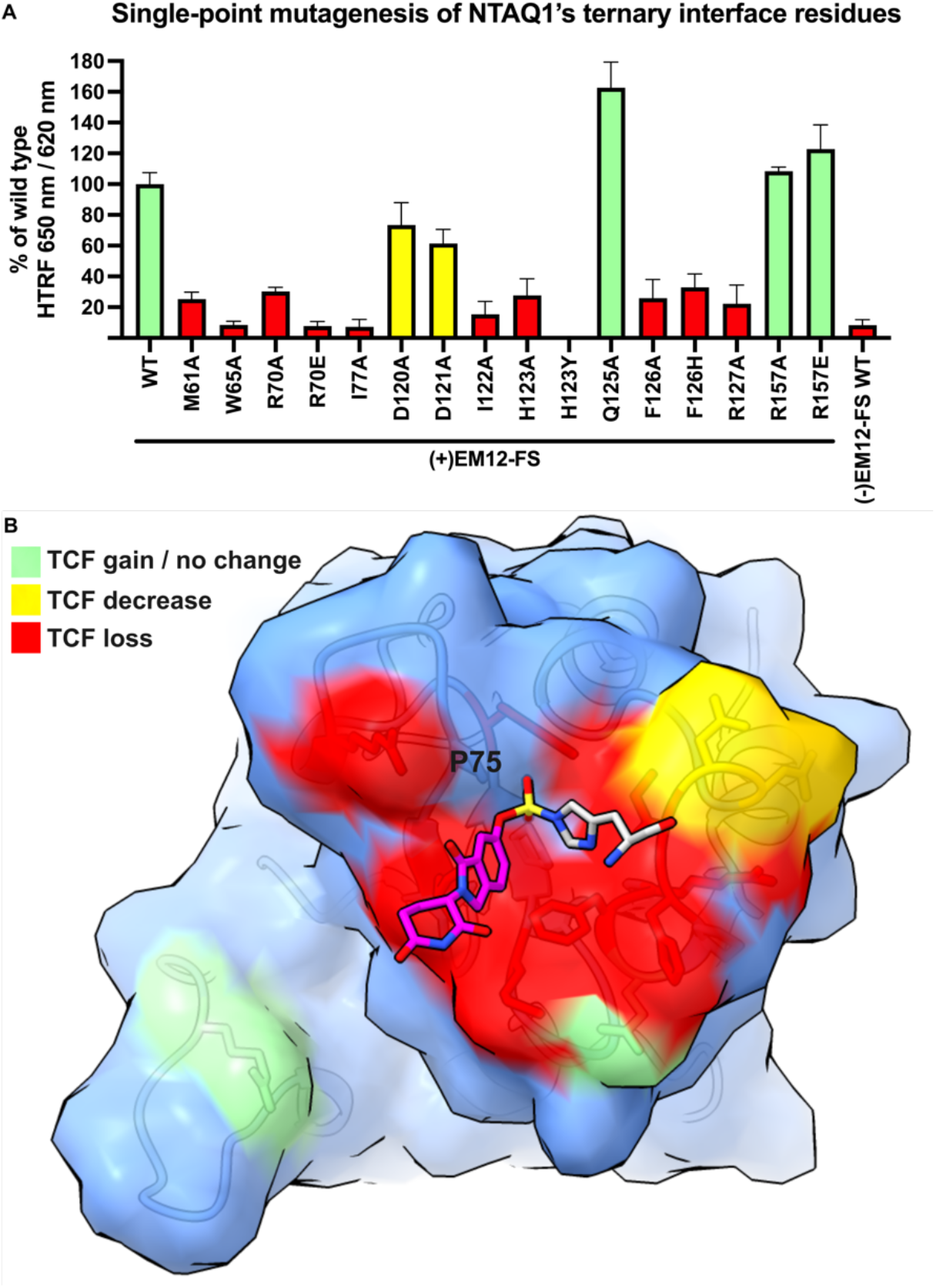
NTAQ1 mutation effects on ternary complex formation (TCF). **A)** TCF HTRF measurements for each NTAQ1 mutant relative to wild-type NTAQ1. Bar colors correspond to mutations that increased (green), decreased (yellow), or abolished (red) TCF. **B)** NTAQ1 PPI surface (blue) as viewed from the CRBN PPI. Mutated residues are colored on the NTAQ1 surface with the same color scheme as in A). EM12-FS (magenta) is shown covalently bound to His353 (gray).

## Discussion

NTAQ1 and the Arg/N-degron pathway have remained underexplored due to the absence of chemical tools.^13^ EM12-FS now provides the first probe for selectively targeting NTAQ1, enabling mechanistic studies of its roles in DNA repair and cancer^17,18^ and supporting structure-guided development of more potent degraders. More broadly, this work reveals the plasticity of the CRBN surface and demonstrates that covalent neofunctionalization can unlock neosubstrate classes inaccessible to reversible IMiDs. By enforcing a closed sensor-loop conformation through covalent modification of His353, EM12-FS creates a non-canonical interface that supports recruitment of a substrate lacking the β-hairpin G-loop degron, establishing a new mode of cereblon engagement.

These findings highlight covalent remodeling as a generalizable strategy for expanding induced-proximity pharmacology. Synthetic modification of E3 ligase surfaces, and potentially other enzyme scaffolds, can reengineer protein-protein interfaces, redirect substrate specificity, and rewire signaling pathways. The requirement for precise electrophile geometry underscores an emerging design principle for covalent molecular glues: the warhead must not only react but must impose a productive conformational state compatible with the intended substrate. Together, these data show that covalent remodeling of CRBN enables synthetic rewiring of induced-proximity interactions and provides a framework for engineering new degrader modalities.

## Acknowledgements

We thank Deerfield for funding.

## Author contributions

A.H.P. and J.T.C. prepared cryo-EM samples, collected and processed cryo-EM data, and performed biochemical experiments. J.C. performed computational studies. L.H.J. conceived and directed the study. L.H.J. wrote the manuscript with input from all authors. L.H.J., M.E.M., P.P.C., and E.S.F. supervised the project.

## Declaration of interests

L.H.J. is a co-founder of Anvia, and scientific advisory board (SAB) member of Anvia, Proxima, Rapafusyn, and Viraccio, an advisor to Flagship, and holds equity in Rapafusyn and Hyku. E.S.F. is a founder, scientific advisory board (SAB) member, and equity holder of Civetta Therapeutics, Proximity Therapeutics, Anvia Therapeutics (also board of directors), Nias Bio, Stelexis Biosciences, Vasetto Bio (also board of directors), HiddenSee Therapeutics, and Neomorph (also board of directors). He is an equity holder and SAB member for Photys Therapeutics and Ajax Therapeutics, and an equity holder in Lighthorse Therapeutics, Sequome and Avilar. E.S.F. is a consultant to Novartis, Eli Lilly and Deerfield. The Fischer lab receives or has received research funding from Deerfield, Novartis, Ajax, Interline, Bayer, and Astellas. A.H.P., M.E.M., and P.P.C. are employees and equity holders in Neomorph and P.P.C. is also a member of the board of directors of Neomorph.

## Data availability

All data supporting the findings of this study are available within the paper and its Supplementary Information. Cryo-EM maps have been deposited in the Electron Microscopy Data Bank under accession number EMD-77682, and the corresponding atomic coordinates have been deposited in the Protein Data Bank under accession number 36MP.

## Supplementary Information

### Supplementary Methods

#### Cloning, expression, and purification of CRBN_Δ1-40_/DDB1_ΔBPB_ for structural studies

Human CRBN_Δ1-40_ and DDB1_ΔBPB_ were cloned into pAC-derived vectors. The recombinant protein complex was co-expressed as StrepTag-Avi-TEV-CRBN_Δ1-40_ and 6His-DDB1_ΔBPB_ in Trichoplusia High-Five insect cells using the baculovirus expression system (Invitrogen). Cells were lysed by sonication in buffer containing 50 mM Tris-HCl pH 8.0, 200 mM NaCl, 1 mM TCEP, 1 mM PMSF and 1x protease inhibitor cocktail (Sigma). Following ultracentrifugation and filtration, the soluble fraction was incubated with Strep-Tactin XT Superflow high-capacity resin (IBA Lifesciences) for 1 hr at 4°C and eluted with buffer containing 100 mM biotin. The complex was cleaved with TEV protease overnight at 4°C before further purification by anion exchange chromatography (Poros 50HQ). Purification was finished by size exclusion chromatography using a HiLoad 16/600 Superdex 200 pg column (Cytiva Lifesciences) in 25 mM HEPES pH 7.4, 200 mM NaCl and 1 mM TCEP. Fractions containing pure TEV-cleaved CRBN_Δ1-40_/DDB1_ΔBPB_ complex were pooled and concentrated using centrifugal ultrafiltration (Millipore), flash frozen in liquid nitrogen, and stored at −80°C.

#### Covalent modification of CRBN/DDB1 with EM12-FS for structural studies

CRBN_Δ1-40_/DDB1_ΔBPB_ was reacted with a 2-fold molar excess of EM12-FS for 24 hours at room temperature. Excess EM12-FS was removed using Zeba Spin Desalting columns (ThermoFischer Scientific) equilibrated with buffer containing 25 mM HEPES pH 7.4, 200 mM NaCl and 1 mM TCEP. Covalently modified CRBN_Δ1-40_/DDB1_ΔBPB_ was flash frozen in liquid nitrogen, and stored at −80°C.

#### Cloning, expression, and purification of NTAQ1 for structural studies

NTAQ1 was purified as previously described. Human wild-type NTAQ1 (Uniprot ID: Q96HA8 isoform 1) was cloned into a pET28a(+) vector and expression of 6His-Avi-TEV-NTAQ1 was induced with the addition of 0.4 mM IPTG in BL-21 Rosetta 2 pLysS *Escherichia coli* cells (Novagen) for 3 hours at 37 °C. Pelleted cells were lysed by sonication in buffer containing 50 mM Tris pH 8.0, 200 mM NaCl, 20 mM imidazole, 1 mM TCEP, and 1 mM PMSF. The lysate was cleared by centrifugation and filtration before incubation with Ni-NTA sepharose (Cytiva) for 1 hour at 4 °C. The protein was eluted with buffer containing 300 mM imidazole. Fractions containing NTAQ1 were pooled and further purified via size-exclusion chromatography using a HiLoad 16/600 Superdex 75 pg column (Cytiva Lifesciences) in 25 mM HEPES pH 7.4, 200 mM NaCl and 1 mM TCEP. Fractions containing NTAQ1 were pooled and concentrated using centrifugal ultrafiltration (Millipore), flash frozen in liquid nitrogen, and stored at −80°C.

#### Cloning, expression, and purification of CRBN/DDB1_ΔBPB_ for biochemical studies

CRBN/DDB1 for TR-FRET was purified as previously described. Human CRBN and DDB1 were cloned into pAC-derived vectors and recombinant protein complex was co-expressed as Flag-Spytag-TEV-CRBN and 6His-Spytag-DDB1_ΔBPB_ in Trichoplusia High-Five insect cells using the baculovirus expression system (Invitrogen). Cells were lysed by sonication in 50 mM Tris-HCl pH 8.0, 200 mM NaCl, 1 mM TCEP, 1 mM PMSF and 1x protease inhibitor cocktail (Sigma). Following ultracentrifugation and filtration, the soluble fraction was incubated with anti-DYKDDDDK resin (Genescript) for 1 hour at 4°C and eluted with buffer containing 150 ug/ml 3X-DYKDDDDK peptide. The complex was further purified by anion exchange chromatography (Poros 50HQ). Following overnight incubation at 4°C with Alexa Fluor 647-C2-labelled SpyCatcher_S50C_, the labeled complex was purified by size exclusion chromatography in 25 mM HEPES pH 7.4, 200 mM NaCl and 1 mM TCEP. Fractions containing labelled CRBN-DDB1 complex were pooled and concentrated using ultrafiltration (Millipore), flash frozen in liquid nitrogen, and stored at −80°C.

#### Cloning, expression, and purification of NTAQ1 and single-point NTAQ1 mutants for biochemical studies

A protein sequence corresponding to NTAQ1 (UniprotKB: Q96HA8-1) with an N-terminal sequence containing a hexahistidine tag, a hemaglutining tag, and a TEV cleavage site were synthesized into the corresponding *E. coli* codon-optimized CDNA. The CDNA was inserted after the Factor Xa site in frame with the maltose binding protein of the pMAL-c5x plasmid (GenScript USA, Inc.). The resulting pMAL-c5x plasmid containing MBP-Xa-6xHis-HA-TEV-NTAQ1 was used as the template to generate single-point mutants of the residues at the NTAQ1 interface with CRBN and EM12-FS.

*E. coli* OneShot BL21(DE3) chemically competent cells (Invitrogen #440048) were transformed with 200 ng of plasmid using a 42 °C heat shock in a water bath followed by 2 minute incubation on ice and 1 hour recovery in SOC medium. The recovered cells were then cultured overnight at 37 °C in 2XYT medium supplemented with 100 ug/mL carbenicillin in a volume of 5 mL. The following morning, 50 mL of 2XYT medium supplemented with 100 ug/mL carbenicillin were inoculated with 1 mL of overnight culture at 37 °C until the cultures reached an optical density (λ=600 nm) of 0.60 and transferred to 18 °C for overnight protein expression. Protein expression was induced by the addition of 250 uM IPTG. The following morning, after approximately 18 hours of growth, the *E. coli* cells were harvested by ultracentrifugation at 4000g using a swinging bucket centrifuge (Beckman Coulter: Avanti J-15R IVD) equilibrated at 4 °C. The cell pellet was stored at -80 °C to await protein purification.

Cell pellets were thawed on ice and resuspended in a solution consisting of 50 mM Tris pH 7.4, 500 mM NaCl, 1 mM TCEP, 5% glycerol, and a 12 U/ml Pierce universal nuclease (TFS #88701). The resuspension was then lyzed by sonication using an ultrasonic processor (U.S. SOLID #JFHLUH00016) operated at 4 °C with 12x 60% power 1-second pulses over 50 seconds. The lysate was clarified in the swinging bucket centrifuge and at 4000g for 30 minutes. The clarified lysate was incubated with 1 ml of amylose resin (BioLabs #E8021L) equilibrated in lysis buffer and incubated in tube rotator (BOEKEL #260750) for 1 hour at 4 °C. The soluble fraction was decanted into a gravity column (BioRad #7321010) and the amylose resin allowed to settle in the column. The column was then washed with 30 ml of a solution consisting of 50 mM Tris pH 7.4, 250 mM NaCl, 1 mM TCEP, 5% glycerol. The washed resin was incubated for 30 minutes with 2 ml of wash buffer supplemented with 10 mM maltose (Sigma #M5895), eluted, syringe filtered with a .22 um filter (Millipore #SLGPR33RS) and concentrated to ∼10 mg/ml using a 10,000 MWCO centrifugal device (Sartorius #VSO4TO2). The resulting protein was analyzed by SDS PAGE (Novex #XP04202BOX, #LC2676, #LC26755 and Abcam #ab119211), aliquoted, flash-frozen in liquid nitrogen, and stored at -80 °C to await biochemical characterization.

#### Homogeneous time-resolved fluorescence (HTRF)

A solution containing 10 nM of CRBN/BBD1 (labeled with Alexa647 and EM12-FS), 20 nM anti-HA antibody (Europium labeled), and PPI-Europium detection buffer (Revvity #61DB9RDF) was made (Solution A) and plated into a black low-volume flat-bottom plate (Corning #3820) with 25 uL per well. Solutions for each of the purified NTAQ1 recombinant proteins were created at a concentration of 2 uM in a buffer containing 20 mM Tris pH 7.5, 100 mM NaCl, and 0.3 % Tween 20). Each of the NTAQ1 solutions was used to plate a 6-point dose response curve in triplicate over the plates containing Solution A using a robotic dispenser (Tecan D300e). The plates were sealed, shaken for 10 minutes, and centrifuged for 5 minutes at 100g using a swinging bucket centrifuge (Beckman Coulter: Avanti J-15R IVD). The plates were allowed to incubate at 4 °C for 1 hour prior to being imaged on a plate reader (PHERAstar FSX) equipped with an HTRF filter module (version F) with a 337 nm excitation, 664 nm emission (channel A) and a 620 nm emission (channel B) using an 80 us delay, 100 flashes, and 270 µs integration time. Each dose response curve was normalize to the per-plate maximum NTAQ1 wild-type response. The HTRF ratio at 20 nM substrate was used for comparison to NTAQ1 wild type since a hook effect was observed at concentrations of substrate exceeding the anti-HA antibody.

#### In vitro ubiquitination and immunoblotting

A ubiquitination system was reconstituted *in vitro* as previously described (DOI:1038/s41589-018-0129-x) with adjustments to allow for the covalent labeling of CRBN.

Two E3 enzyme CRBN-DDB1-Cul4-Rbx1-Nedd8 mixtures were prepared ahead of time to allow for 24-hour treatment with DMSO or EM12-FS (unlabeled-E3 or labeled-E3, respectively). To prepare unlabeled-E3, 1.1 uL of 0.5 % DMSO was added to 56.2 uL of 2.4 uM CRBN-DDB1 and allowed to incubate for 24 hrs at room temperature in 20 mM HEPES pH 7.5, 150 mM NaCl. Following incubation, 8.8 uL of 15.5 uM Cul4-Rbx1-Nedd8 was added to obtain a final concentration of ∼2 uM unlabeled-E3. To prepare labeled-E3, 1.1 uL of 500 uM EM12-FS was added to 56.2 uL of 2.4 uM CRBN-DDB1 and allowed to incubate for 24 hrs at room temperature in 20 mM HEPES pH 7.5, 150 mM NaCl. Following incubation, 8.8 uL of 15.5 uM Cul4-Rbx1-Nedd8 was added to obtain a concentration of ∼2 uM labeled-E3.

An E1-E2 mixture containing 200 nM Ube1, 1 uM UbcH5a, 1 uM UbcH5b, 200 uM Ubiquitin, and 40 mM ATP was prepared in 20 mM HEPES pH 7.5, 150 mM NaCl_2_, 10 mM MgCl_2_. Additionally, 30 uM substrate stocks of WT NTAQ1 and single-point mutant NTAQ1 were prepared by diluting the previously purified recombinant proteins.

To conduct the ubiquitination reactions, 11.8 uL of E1-E2 mix, 6.5 uL of E3 mix (labeled or unlabeled), and 5 uL of NTAQ1 substrate were combined and incubated for 2.5 hours at 30 °C. The reactions were then stopped by the addition of 25 uL 2x SDS-PAGE running buffer and incubation at 95 °C for 2 minutes. The samples were then separated by SDS-PAGE in preparation for Coomassie staining and immunoblotting. Immunoblotting was conducted using standard Western blot protocols with a Trans-Blot Turbo transfer system in high molecular weight mode, 1:1000 mouse anti-MBP (NEB: E8032S) primary antibody incubation at 4 °C overnight, 1:20000 goat anti-mouse (LI-COR: 926-32210) secondary antibody incubation at room temperature for 1 hour, and exposure with a LI-COR Odyssey CLx imaging system.

#### Grid preparation for cryo-electron microscopy

A solution containing 10 uM of CRBN/DDB1dBPB, pre-labeled with EM12-FS, and 15 uM of NTAQ1 was incubated on ice for 15 minutes before a 3-fold dilution into a solution containing 10 mM HEPES pH 7.4, 240 mM NaCl, 1 mM TCEP, and 0.01% lauryl maltose neopentyl glycol. 4 uL of the diluent were applied to a Quantifoil 300 mesh R1.2/1.3 UltrAuFoil (EMS #Q350AR13A) grid that had been treated with a PELCO EasiGlow glow discharge cleaning system (TED PELLA #91000) and preloaded onto a Vitrobot Mark IV system (TFS #1151972) equilibrated at 95% humidity and 4 °C. Following application of the diluent, the grid was blotted for 4 seconds by the Vitrobot using standard blotting paper (EMS #71166-65) and immediately plunged into liquid ethane for vitrification. The vitrified grid was stored under cryogenic conditions to await data acquisition.

#### Cryo-electron microscopy data acquisition and single particle analysis

Cryo-EM data were acquired using the Gatan DigitalMicrograph software and a Titan Krios G3 equipped with a Gatan K3 direct electron detector and a Gatan Biocontinuum Energy Filter. Movies were recorded at a magnification of 81,000x (magnified pixel size of 1.072 Å) with a total dose of 41 e^-^/A^2^ over 5.5 seconds.

#### Cryo-electron microscopy data processing

Frame alignment, dose weighting, CTF estimation, particle extraction, and 2D class averaging were performed in real-time using cryoSparc v4.4. An initial model with a resolution of 15 Å was generated from an atomic model of CRBN/DDB1dBPB (no substrate) in Chimera 1.17 using molmap. Extensive 3D classification was performed in cryoSparc and Relion 5.0-beta to obtain a homogeneous subset of particles with an anisotropic angular distribution. CTF refinement, gold-standard FSC calculations, and local resolution estimation were performed in Relion. The final reconstruction was then sharpened using Relion PostProcess and DeepEMhancer.

#### Atomic model building and deposition

Alphafold models for DDB1 (AF-Q16531-F1), CRBN (AF-Q96SW2-F1), and NTAQ1 (AF-Q96HA8-F1) were rigid body docked into the sharpened map from DeepEMhancer using Chimera (full domain) and Coot 0.9 (subdomains). Propeller B of DDB1 was removed in Coot and the atomic model was real-spaced refined in Phenix 1.21-5207. Several cycles of automatic refinement (Phenix) and manual refinement (Coot) were performed. A custom crystallographic information file (CIF) dictionary was generated for CRBN’s histidine 353 covalently bound to EM12-FS using the eLBOW module in Phenix. This modified histidine was used throughout the atomic model refinement process. Following model building and refinement with the DeepEMhancer map, the atomic model was refined with the Relion sharpened map for deposition. The atomic model was deposited in the RCSB Protein Data Bank with accession code PDB: 36MP.

#### Molecular modeling and MM-GBSA

EM12-SF-CRBN was modeled based on the cryo-EM structure with EM12-FS. EM12-SF was first aligned to EM12-FS using the substructure EM12, the covalent bond between sulfone and His353 was then built manually. The ternary complex structure was then optimized using the “Refine Receptor-Ligand complex” protocol in Schrodinger suite (2025-1) with default parameters. The MM-GBSA calculations were performed by defining NTAQ1 as the ligand. Residues within 4 Å of the ligand (i.e. NTAQ1) were considered flexible. The affinity difference estimated by MM-GBSA was 6 kcal/mol in favor of EM12-FS.

**Figure S1.**
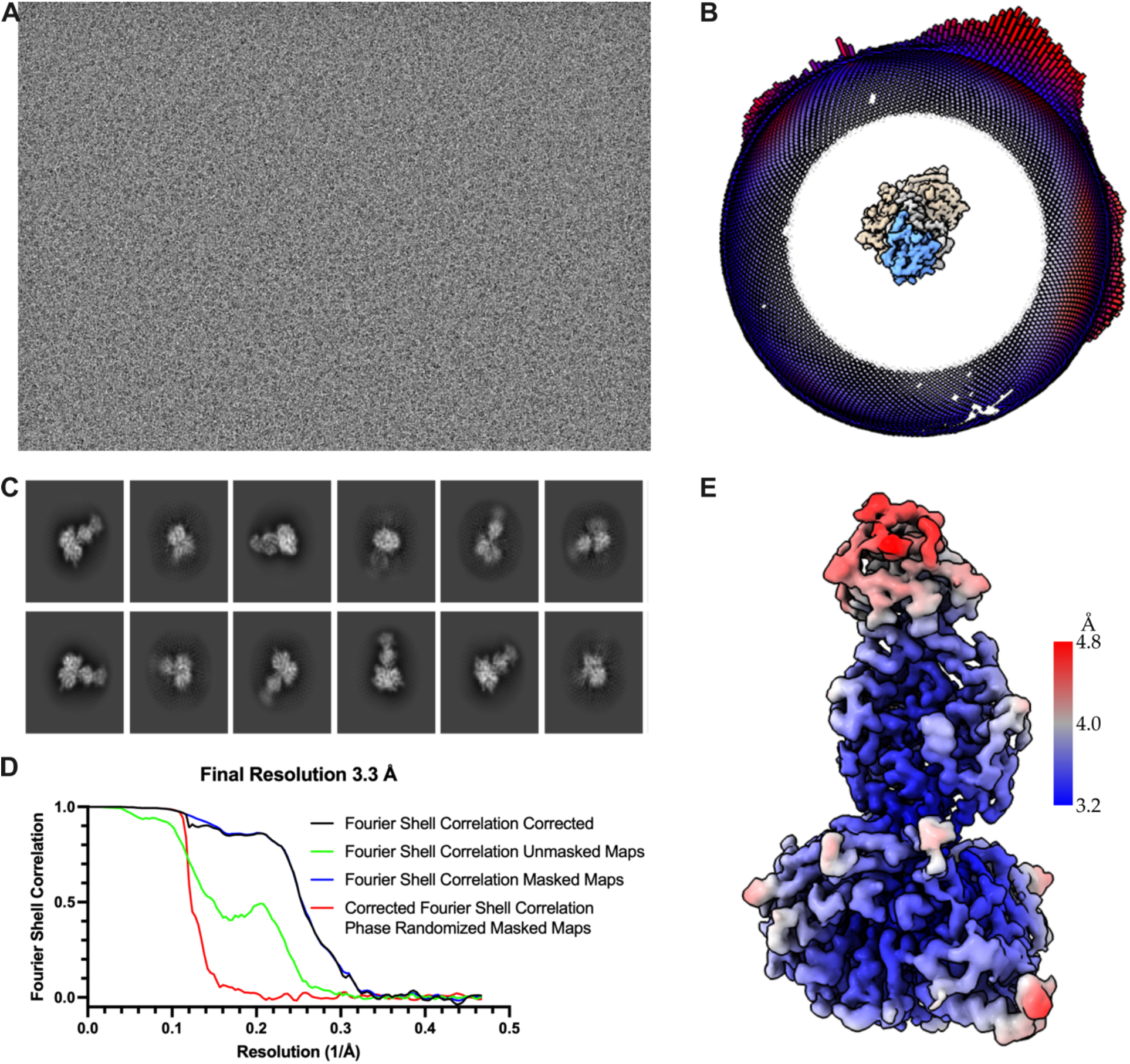
Cryo-EM single-particle analysis. **A)** Representative micrograph. **B)** Single-particle angular distributions showing more abundant views as larger bars (red) and less abundant views as smaller bars (blue). The ternary complex is shown for reference with NTAQ1 in light blue, CRBN in gray, and DDB1 in tan. **C)** Representative 2D class averages. **D)** Gold-standard FSC calculated in RELION for the final cryo-EM reconstruction. **E**. Local resolution estimate of the final cryo-EM reconstruction.

**Figure S2.**
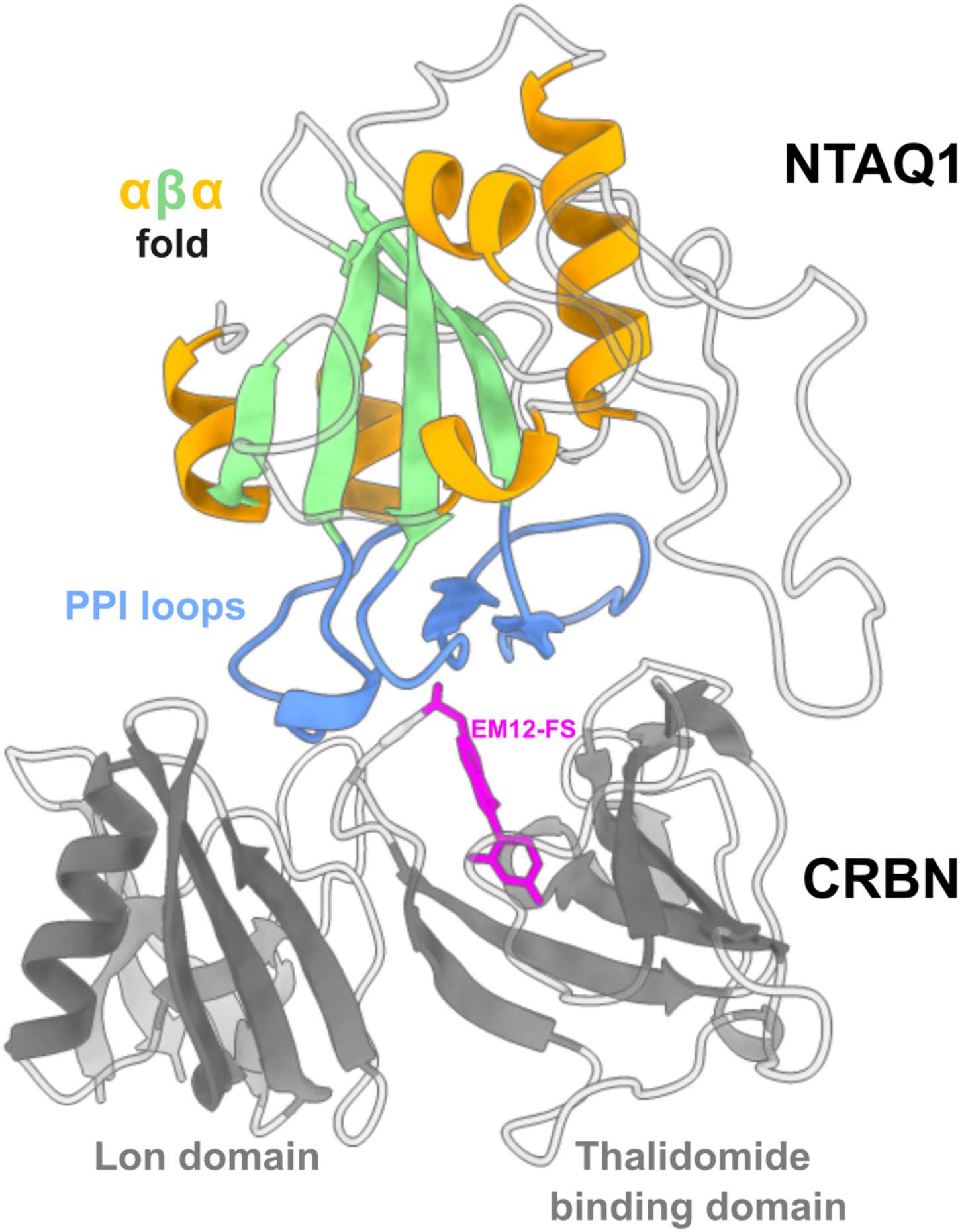
Ternary complex overview. NTAQ1 recruitment to CRBN is shown with the NTAQ1 elements primarily forming the PPI interface colored in blue. The NTAQ1 αβα fold is shown with α-helices and green β-sheets. CRBN is shown in gray and EM12-FS in magenta. Disordered loops are shown uncolored.

**Figure S3.**
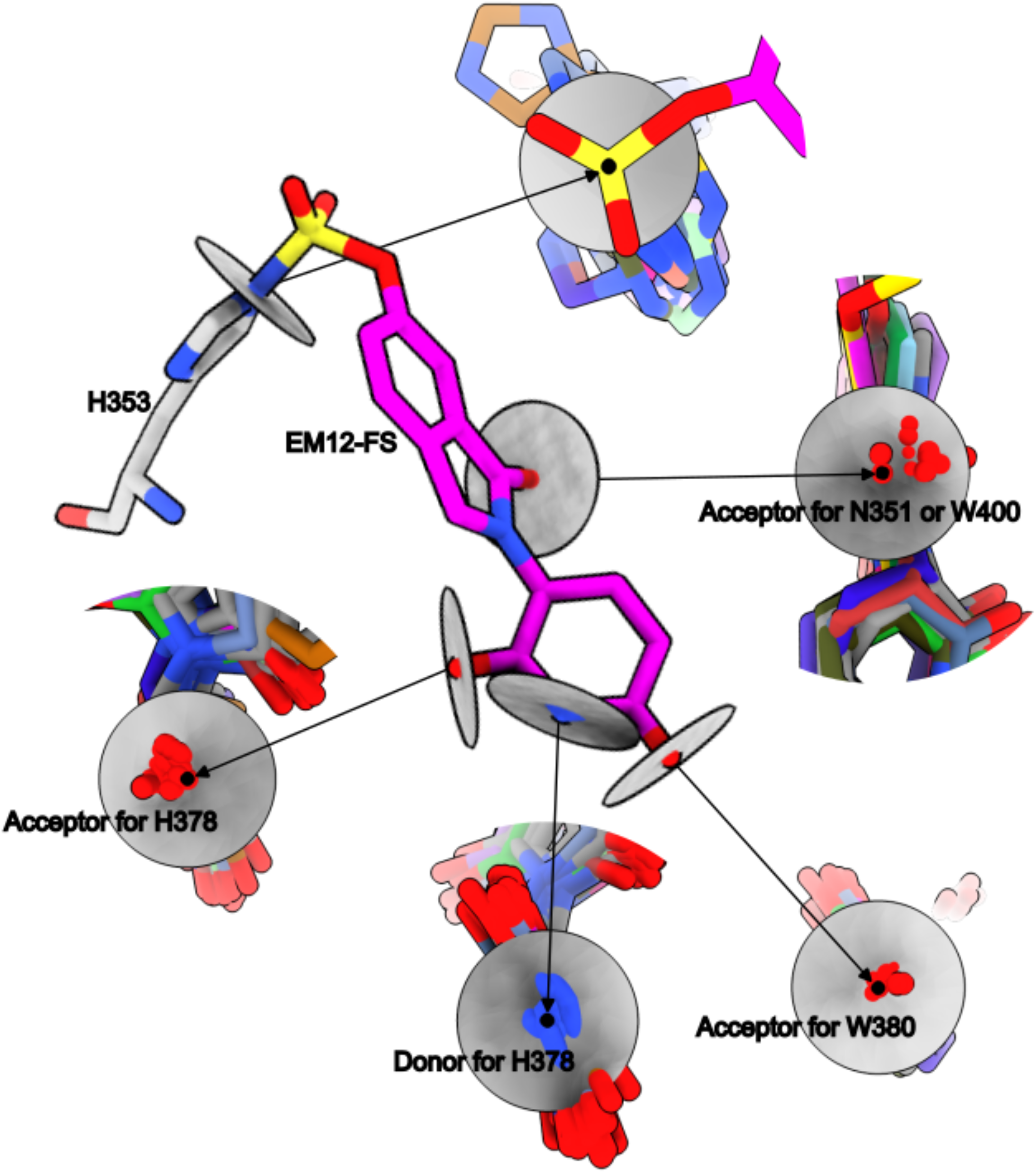
EM12-FS binding mode comparison to non-covalent small molecule human CRBN binders deposited in the RCSB PDB. Human CRBN structures below 3.5 Å were download from the RCSB PDB and aligned to the tri-tryptophan pocket of the EM12-FS ternary complex structure (PDB IDs: 4CI1, 4CI2, 4CI3, 4TZ4, 5FQD, 5V3O, 6BOY, 6H0F, 7BQU, 7BQV, 7U8F, 8D7U, 8D7V, 8D7W, 8D7Z, 8G66, 8OIZ, 8OJH, 8RQ8, 8RQ9, 8RQA, 8RQC, 8TNP, 8TNQ, 8TNR, 8TZX, 8U15, 8U16, 8U17, 9CUO, 9DJX, 9DQD, 9FJX). The protein was hidden (except for Histidine 353) and the compounds were colored differentially at their carbons. EM12-FS was colored in magenta. A plane (gray) normal to the indicated carbonyls (red) and nitrogens (blue) is shown to facilitate visualization of the relative orientation of other CRBN binders to EM12-FS (center of plane, black dot). A plane is also shown for Histidine 353 on CRBN to show the differences observed in the EM12-FS ternary complex.

**Figure S4.**
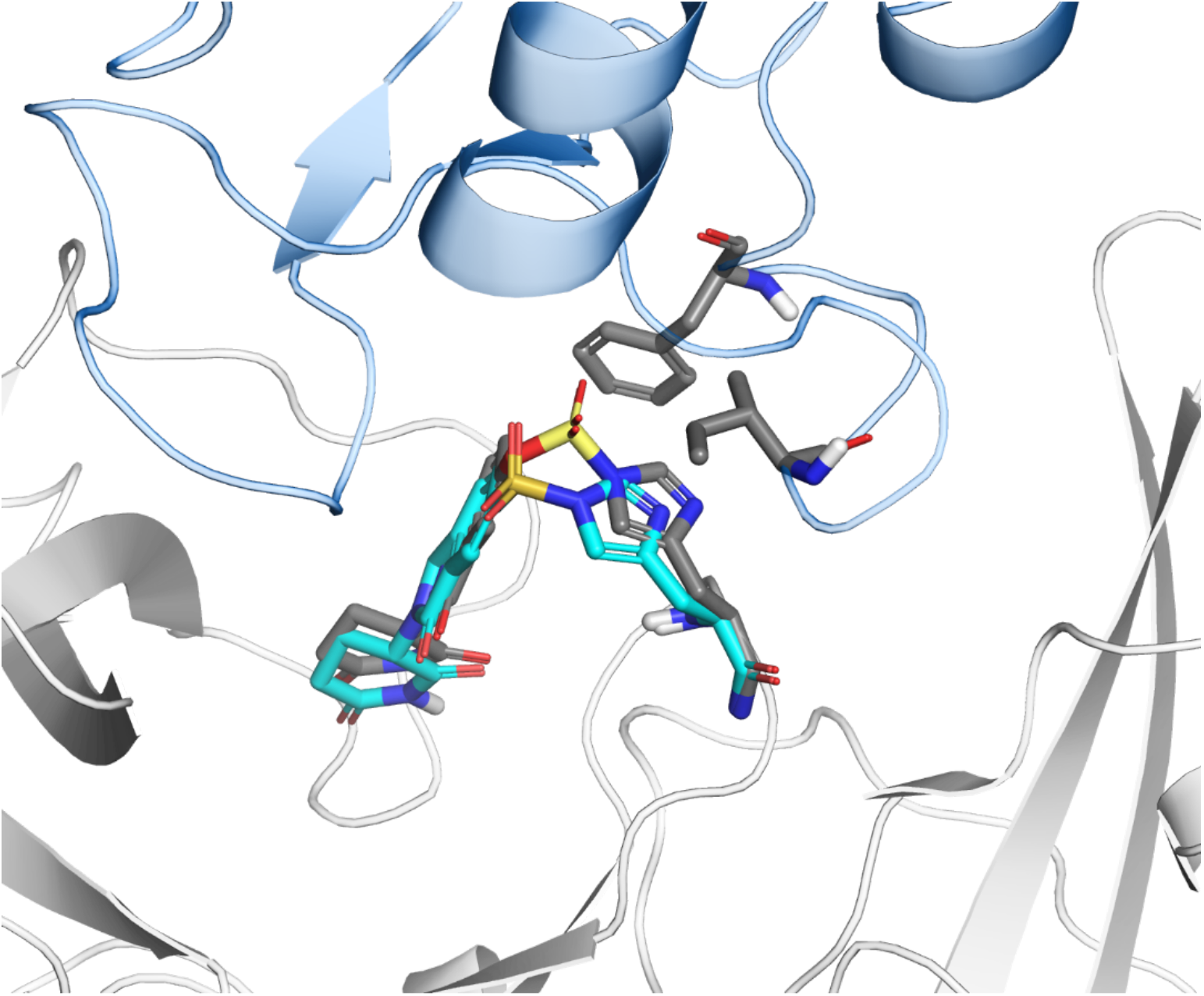
Molecular model of EM12-SF–CRBN (EM12-SF/His353 in cyan) overlayed with the EM12-FS cryo-EM structure (EM12-FS/His353 in gray). The imidazole ring of His353 is pulled further away from NTAQ1 residues Ile122 and Phe126 (gray) by EM12-SF.

**Figure S5.**
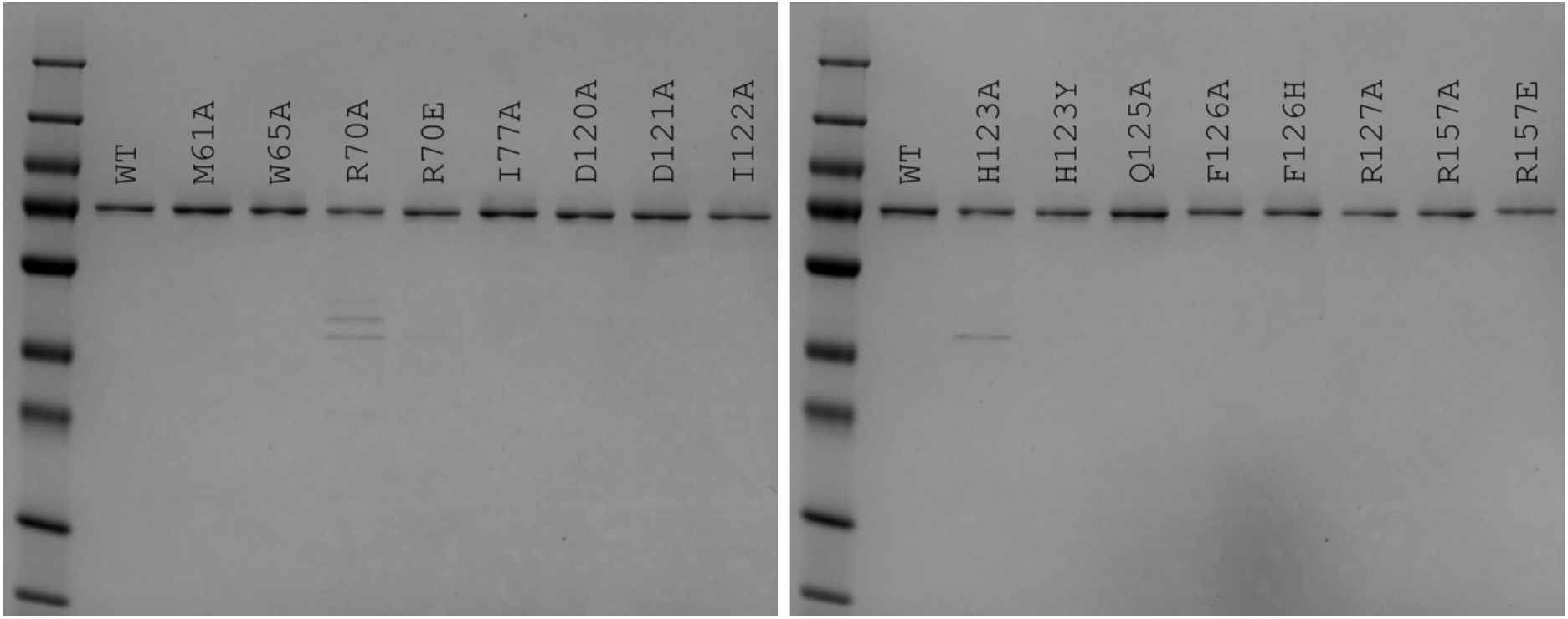
SDS-PAGE of NTAQ1recombinant protein constructs purified for biochemical studies.

**Figure S6.**
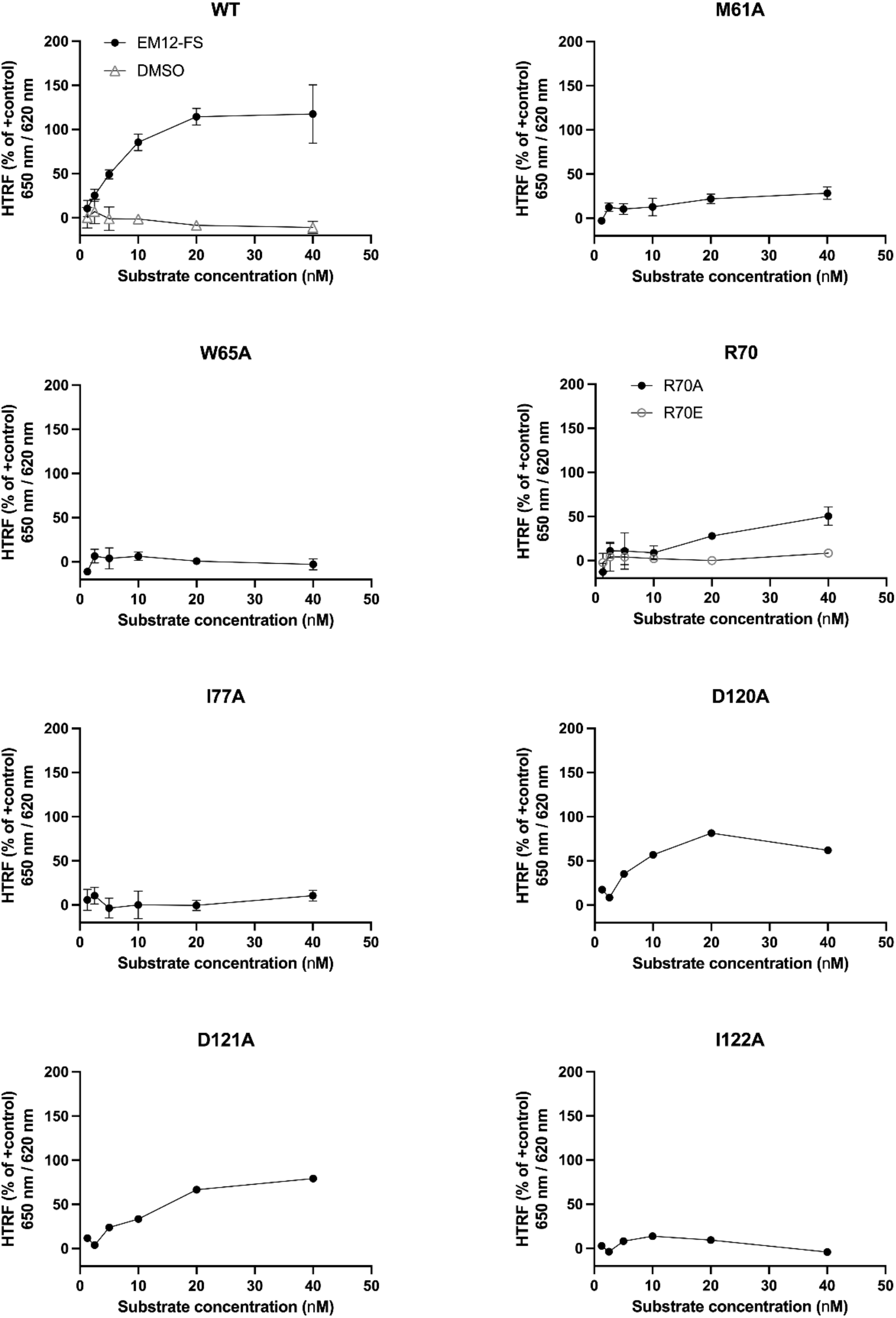
Ternary complex formation homogeneous time-resolved fluorescence of NTAQ1 wild type or single-point mutants with CRBN. The assayed NTAQ1 protein (WT or mutant) is indicated above each panel. Wild type CRBN was incubated with EM12-FS (circle markers) or DMSO (triangle markers, top-left panel). For NTAQ1 residues with more than one mutation type, the additional mutant is shown with an empty circle marker.

**Figure S7.**
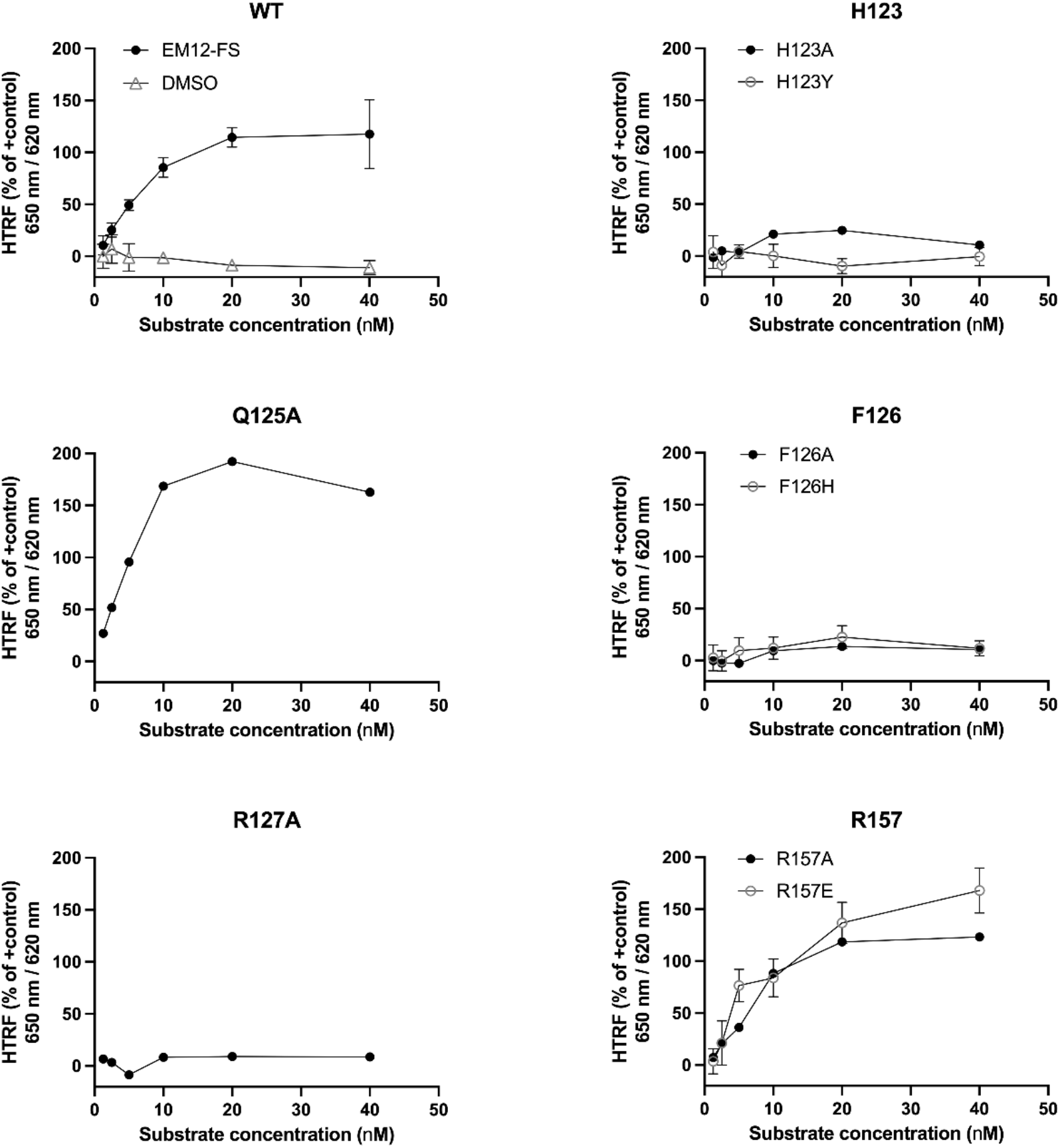
Ternary complex formation homogeneous time-resolved fluorescence of NTAQ1 wild type or single-point mutants with CRBN. The assayed NTAQ1 protein (WT or mutant) is indicated above each panel. Wild type CRBN was incubated with EM12-FS (circle markers) or DMSO (triangle markers, top-left panel). For NTAQ1 residues with more than one mutation type, the additional mutant is shown with an empty circle marker.

**Figure S8.**
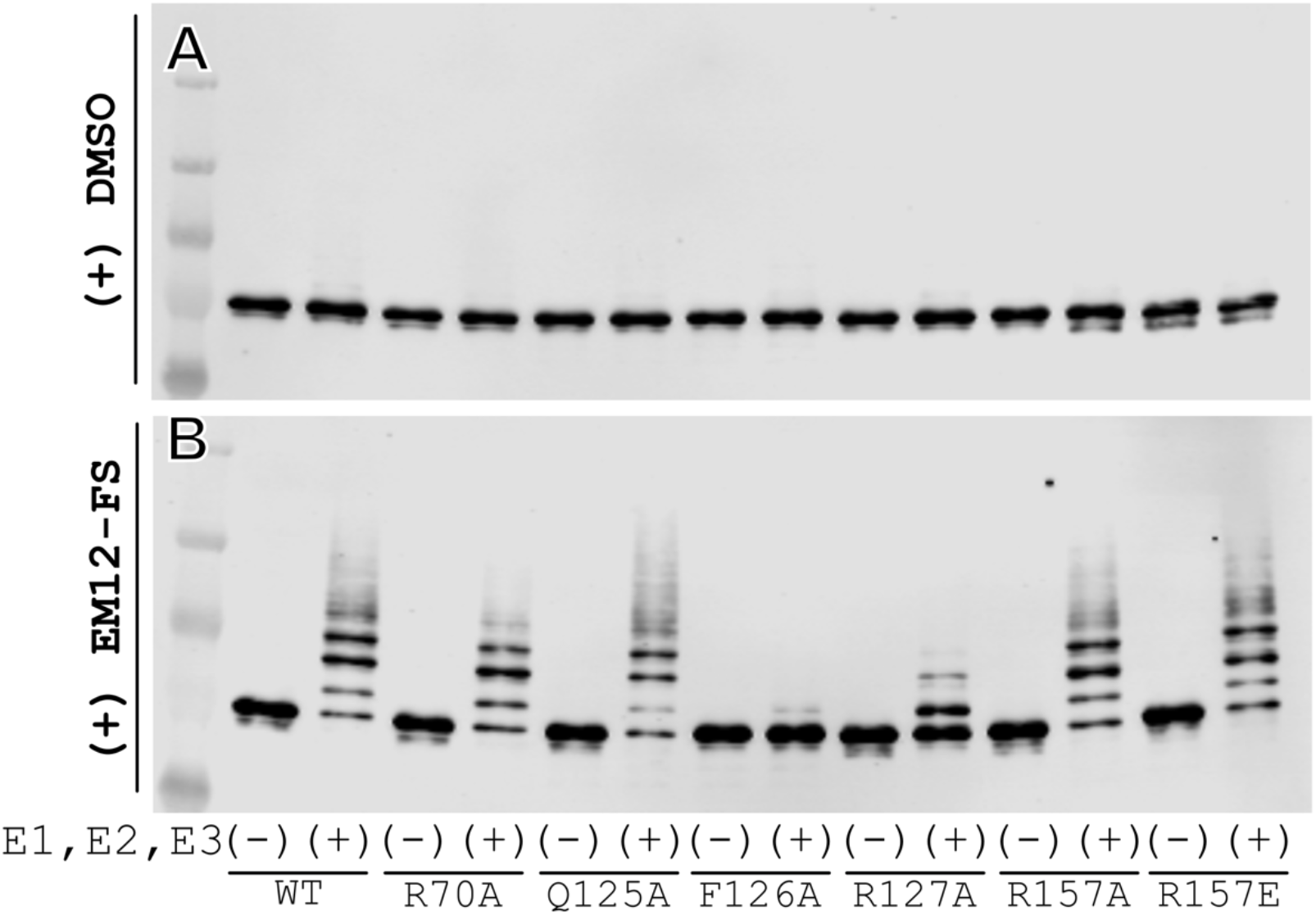
In vitro ubiquitination immunoblot. NTAQ1 WT and select single-point mutants were assayed with a reconstituted in vitro ubiquitination system in the presence of CRBN which had been treated with DMSO (Panel A) or EM12-FS (Panel B). Each protein construct was tested with (+) and without (-) the reconstituted system, separated by SDS-PAGE, and visualized by Western blot using an antibody against the recombinant MBP tag on each protein construct.

**Table S1.** CRBN-NTAQ1 interactions.

| CRBN residue | NTAQ1 residue | Type of interaction | Distance |
| --- | --- | --- | --- |
| His353 | Phe126 | edge-to-face π-interaction | 4.2 Å |
| His353 | His123 (backbone) | hydrogen bond | 4.2 Å |
| EM12-FS | Trp65 | hydrogen bond | 4.6 Å |
| His103 | Asp121 | hydrogen bond | 2.6 Å |
| Asp149 | Gln125 | hydrogen bond | 4.0 Å |
| Glu377 | Met61 | hydrophobic | 4.8 Å |
| Gln390 | Arg70 | hydrogen bond | 2.5 Å |
| Gln390 | Pro71 (backbone) | hydrogen bond | 3.6 Å |
| Phe102 | Arg127 | cation-π interaction | 4.8 Å |
| Trp386 | Ile77 | hydrophobic | 4.8 Å |
| Ile152 and 154 | Pro124 | hydrophobic | 4.1 and 3.6 Å |
| Phe102 | His123 | hydrophobic | 3.6 Å |
| Pro352 | Phe126 | hydrophobic | 4.2 Å |

## References

1. Ito, T., Ando, H., Suzuki, T., Ogura, T., Hotta, K., Imamura, Y., Yamaguchi, Y., and Handa, H. (2010). Identification of a primary target of thalidomide teratogenicity. Science 327, 1345–1350. 10.1126/science.1177319.

2. Fischer, E.S., Böhm, K., Lydeard, J.R., Yang, H., Stadler, M.B., Cavadini, S., Nagel, J., Serluca, F., Acker, V., Lingaraju, G.M., et al. (2014). Structure of the DDB1-CRBN E3 ubiquitin ligase in complex with thalidomide. Nature 512, 49–53. 10.1038/nature13527.

3. Chamberlain, P.P., Lopez-Girona, A., Miller, K., Carmel, G., Pagarigan, B., Chie-Leon, B., Rychak, E., Corral, L.G., Ren, Y.J., Wang, M., et al. (2014). Structure of the human Cereblon-DDB1-lenalidomide complex reveals basis for responsiveness to thalidomide analogs. Nat Struct Mol Biol 21, 803–809. 10.1038/nsmb.2874.

4. Petzold, G., Fischer, E.S., and Thomä, N.H. (2016). Structural basis of lenalidomide-induced CK1α degradation by the CRL4(CRBN) ubiquitin ligase. Nature 532, 127–130. 10.1038/nature16979.

5. Matyskiela, M.E., Lu, G., Ito, T., Pagarigan, B., Lu, C.C., Miller, K., Fang, W., Wang, N.Y., Nguyen, D., Houston, J., et al. (2016). A novel cereblon modulator recruits GSPT1 to the CRL4(CRBN) ubiquitin ligase. Nature 535, 252–257. 10.1038/nature18611.

6. Sievers, Q.L., Petzold, G., Bunker, R.D., Renneville, A., Słabicki, M., Liddicoat, B.J., Abdulrahman, W., Mikkelsen, T., Ebert, B.L., and Thomä, N.H. (2018). Defining the human C2H2 zinc finger degrome targeted by thalidomide analogs through CRBN. Science 362. 10.1126/science.aat0572.

7. Krönke, J., Fink, E.C., Hollenbach, P.W., MacBeth, K.J., Hurst, S.N., Udeshi, N.D., Chamberlain, P.P., Mani, D.R., Man, H.W., Gandhi, A.K., et al. (2015). Lenalidomide induces ubiquitination and degradation of CK1α in del(5q) MDS. Nature 523, 183–188. 10.1038/nature14610.

8. Petzold, G., Gainza, P., Annunziato, S., Lamberto, I., Trenh, P., McAllister, L.A., DeMarco, B., Schwander, L., Bunker, R.D., Zlotosch, M., et al. (2025). Mining the CRBN target space redefines rules for molecular glue-induced neosubstrate recognition. Science 389, eadt6736. 10.1126/science.adt6736.

9. Steger, M., Nishiguchi, G., Wu, Q., Schwalb, B., Shashikadze, B., McGowan, K., Actis, M., Aggarwal, A., Shi, Z., Price, J., et al. (2025). Unbiased mapping of cereblon neosubstrate landscape by high-throughput proteomics. Nat Commun 16, 7773. 10.1038/s41467-025-62829-0.

10. Annunziato, S., Quan, C., Donckele, E.J., Lamberto, I., Bunker, R.D., Zlotosch, M., Schwander, L., Murthy, A., Wiedmer, L., Staehly, C., et al. (2026). Cereblon induces G3BP2 neosubstrate degradation using molecular surface mimicry. Nat Struct Mol Biol 33, 479–487. 10.1038/s41594-025-01738-8.

11. Ojeda, S., Wang, M., Baek, K., Bourgeois, W., Sommerschield, A., Yue, H., Metivier, R.J., Karagiannis, P., Levitz, T.S., Xiong, Y., et al. (2026). Degron-independent recruitment of KAT2A expands the target space of CRBN molecular glues. Science 393, 188–194. 10.1126/science.aef5391.

12. Jones, L.H. (2024). Synthetic modification of protein surfaces to mediate induced-proximity pharmacology. RSC Med Chem 15, 2974–2979. 10.1039/d4md00388h.

13. Cruite, J.T., Dann, G.P., Che, J., Donovan, K.A., Ferrao, S., Ficarro, S.B., Fischer, E.S., Gray, N.S., Huerta, F., Kong, N.R., et al. (2022). Cereblon covalent modulation through structure-based design of histidine targeting chemical probes. RSC Chem Biol 3, 1105–1110. 10.1039/d2cb00078d.

14. King, E.A., Cho, Y., Hsu, N.S., Dovala, D., McKenna, J.M., Tallarico, J.A., Schirle, M., and Nomura, D.K. (2023). Chemoproteomics-enabled discovery of a covalent molecular glue degrader targeting NF-κB. Cell Chem Biol 30, 394–402.e399. 10.1016/j.chembiol.2023.02.008.

15. Watson, E.R., Novick, S., Matyskiela, M.E., Chamberlain, P.P., H de la Peña, A., Zhu, J., Tran, E., Griffin, P.R., Wertz, I.E., and Lander, G.C. (2022). Molecular glue CELMoD compounds are regulators of cereblon conformation. Science 378, 549–553. 10.1126/science.add7574.

16. Wang, H., Piatkov, K.I., Brower, C.S., and Varshavsky, A. (2009). Glutamine-specific N-terminal amidase, a component of the N-end rule pathway. Mol Cell 34, 686–695. 10.1016/j.molcel.2009.04.032.

17. Piatkov, K.I., Colnaghi, L., Békés, M., Varshavsky, A., and Huang, T.T. (2012). The auto-generated fragment of the Usp1 deubiquitylase is a physiological substrate of the N-end rule pathway. Mol Cell 48, 926–933. 10.1016/j.molcel.2012.10.012.

18. Leboeuf, D., Abakumova, T., Prikazchikova, T., Rhym, L., Anderson, D.G., Zatsepin, T.S., and Piatkov, K.I. (2020). Downregulation of the Arg/N-degron Pathway Sensitizes Cancer Cells to Chemotherapy In Vivo. Mol Ther 28, 1092–1104. 10.1016/j.ymthe.2020.01.021.

